# Structure-Guided Design of a Bioactive Covalent Small Molecule Targeting a Riboswitch

**DOI:** 10.1101/2025.08.01.668122

**Authors:** Chungen Li, Xueyi Yang, Kyle A. Dickerson, Patrick R. A. Zanon, Noah A. Springer, Jielei Wang, Yilin Jia, Nikhil C. Munshi, Robert T. Batey, Matthew D. Disney

## Abstract

Small molecule ligands targeting structured RNA elements hold promise for modulating RNA function, serving as chemical probes and potential therapeutics. In this study, the characterization of phenylglyoxal-based covalent probe designed to target unpaired guanine residues in structured RNAs is reported. A structure-guided design strategy was employed to modify covalently unpaired guanines critical for flavin mononucleotide (FMN) binding to the FMN riboswitch. Covalent modification occurs at the designed site and modulates riboswitch function in a cellular reporter system, highlighting the potential of covalent mechanisms of action for bioactive RNA ligands.

---

RNA is a compelling therapeutic target, and expanding small molecules that modulate RNA function would extend the landscape of drug discovery for human diseases. Targeting RNA transcripts offers an innovative strategy to overcome challenges associated with protein targets, particularly those that are intrinsically disordered or thought to be “undruggable”. Covalent modulators have emerged as a powerful strategy in drug discovery, with proven success for proteins and growing interest for RNA.^1^ The advancement of covalent RNA ligands is constrained by the lack of selective reactive warheads. Several groups have developed covalent RNA modifiers. For example, cellular occupancy and augmentation of the biological activity for small molecules binding RNA has been accomplished by appending *N*-(2-chloroethyl)anilines onto RNA-binding small molecules and has revealed that small molecules can be designed to be specific to a disease-driving allele for RNA repeat expansions.^2, 3^ Additionally other subsequent approaches have used, including diazirines, activated esters, epoxides, α-halo-carbonyls, alkyl bromides and 3-chloropivalamides.^4, 5, 6^

Most of these electrophiles preferentially react with the N7 of guanine residues, the position with the highest intrinsic nu-cleophilicity.^4, 5, 7^ As the N7 of both paired and unpaired guanine residues are reactive, and therefore a large proportion of the transcriptome may be subject to modification, which in some cases could be beneficial to identify binding ligands in cells. However, this broad reactivity could hinder the development of selective covalent RNA ligands. This study focused on identifying and studying alternative reactivities targeting a different and more distinct subset of RNA nucleophiles.

Glyoxal and its derivatives, methylglyoxal, *N*_3_-kethoxal and phenylglyoxal, have been reported to exhibit robust reactivities toward RNA through a distinct covalent mechanism.^8^ In aqueous environments, glyoxal derivatives exist in equilibrium between a hydrated diol and a reactive dicarbonyl form and selectively react with the nucleophilic N1 and N2 atoms at the Watson–Crick faces of unpaired guanines to form two possible imidazopurinone regioisomers (Fig. 1A). Among the glyoxal derivatives, phenylglyoxal offers a moderate, tunable reactivity, favorable cell permeability, and synthetic versatility, and thus a strategic warhead for the design of highly selective, covalent RNA-targeted modulators.^9^

**Figure 1.**
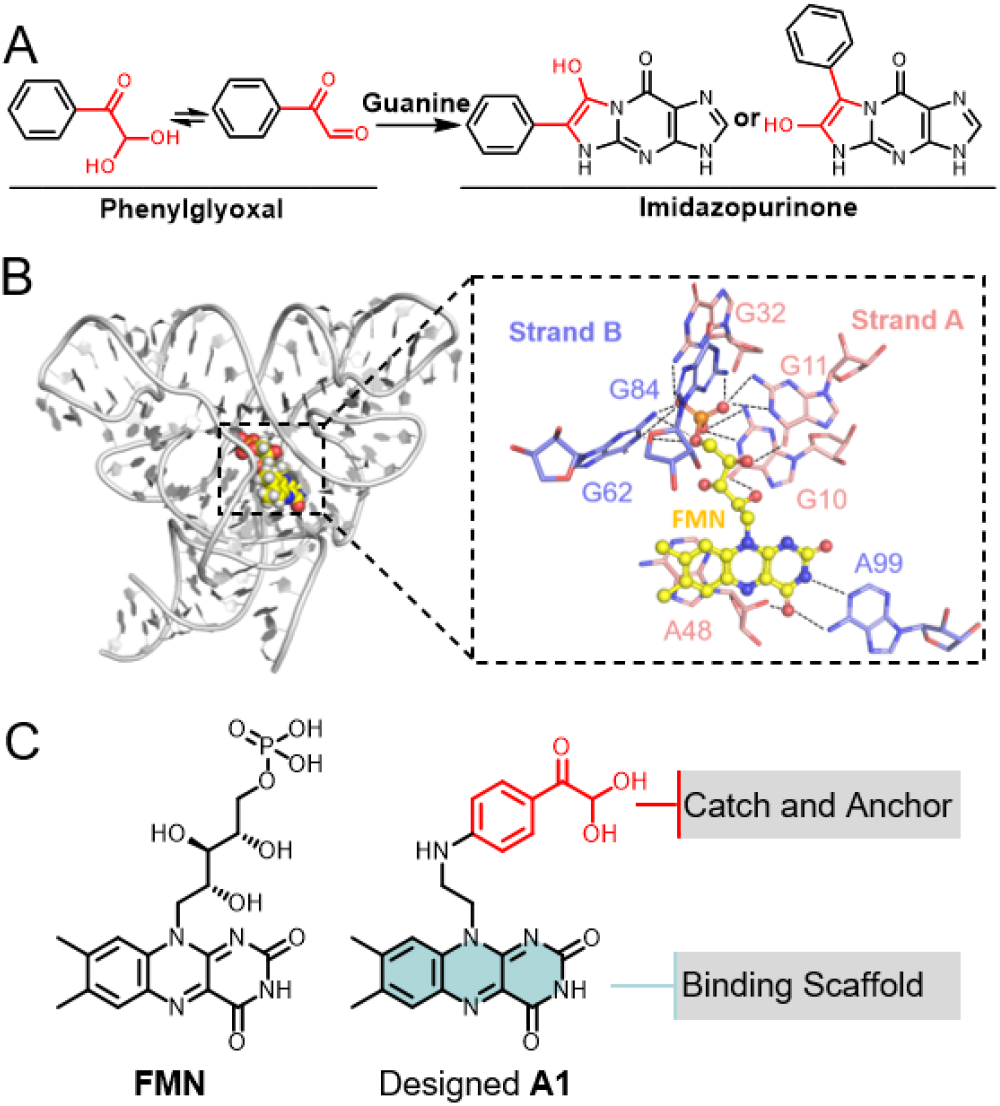
Structure-guided design of a covalent modulator targeting the FMN riboswitch *via* reaction with unpaired guanines. **A)** Reaction of phenylglyoxal with guanine yielding two possible regioisomers. **B)** Crystal structure of the *Fusobacterium nucleatum* FMN riboswitch (gray ribbon) bound to **FMN** (yellow spheres) with stick representation of the FMN binding pocket. The hydrogen bonds are displayed as black dashed lines (PDB ID: 2YIE). **C)** Chemical structures of **FMN** and designed compound **A1**.

The reactivity and selectivity of phenylglyoxal (**P1**) and chlorambucil (**P2**) alkyne probes with r(G_4_C_2_)_8_ and a fully paired RNA r(G_2_C_2_)_8_ were measured using a tetramethylrhodamine (TAMRA) labeling assay (Fig. S1A). The r(G_4_C_2_)_8_ RNA construct harbors six unpaired 1×1 nucleotide GG internal loops embedded in a hairpin, providing multiple potential sites of modification.^10^ In contrast, the r(G_2_C_2_)_8_ RNA construct features sixteen GC base pairs, rendering the Watson–Crick faces of the guanines inaccessible. Following incubation with the RNA, the probes were reacted with a TAMRA-azide dye by a copper catalyzed azide-alkyne cycloaddition (CuAAC) “Click” reaction;^11^ the extent of covalent modification was then measured following gel electrophoresis and fluorescence imaging (Fig. S1B). Compared to the chlorambucil **P2**, the phenylglyoxal **P1** exhibited a milder reactivity against r(G_4_C_2_)_8_ at 200 μM, yielding only 57% of the fluorescence observed with chlorambucil. In contrast, **P1** produced no detectable fluorescence signal with the fully paired r(G_2_C_2_)_8_, indicating a promising selectivity (Fig. S1C, D). Moreover, **P1** reacted dose- and time-dependently with r(G_4_C_2_)_8_, supporting its potential as a warhead for covalent RNA ligands (Fig. S1E, F).

Armed with data to support **P1** as a warhead, we hypothesized that selective covalent modification of an RNA of interest could be achieved by conjugating **P1** to an RNA-binding small molecule. A structure-guided design strategy was employed by analyzing RNA-ligand complex structures in the Protein Data Bank. In particular, we sought to identify RNA targets in which a small molecule binds in proximity to unpaired guanine residues, allowing the phenylglyoxal moiety to react with a specific RNA target *via* binding induced proximity.

The flavin mononucleotide (FMN) riboswitch, an RNA element located in the 5′-leader sequence of bacterial mRNAs that regulates gene expression in response to riboflavin metabolites, has been structurally well characterized and features a ligand-accessible pocket flanked by unpaired guanines.^12, 13^ It was selected for a structure-guided covalent ligand design as a crystal structure revealed that five guanine residues (G10, G11, G32, G62, and G84) form multiple hydrogen bonds with the phosphate moiety of **FMN**. Notably, three of these residues (G10, G11, and G62) are unpaired and possess accessible Watson– Crick faces (Fig. 1B).^14, 15^ We hypothesized that these unpaired guanines could be targeted by replacing the phosphoribosyl of **FMN** with a phenylglyoxal electrophile, affording designed compound **A1** (Fig. 1C).

A non-covalent molecular docking analysis was performed to investigate the potential binding mode of the **A1**. Given that the dehydration product **A1’** is considered the predominantly reactive species, molecular docking was performed with this dicarbonyl species (Fig. S2A). The results suggest **A1’** could bind within the FMN binding pocket and assume a pose like **FMN** (Fig. S2B). The flavin core of **A1’** was predicted to form key hydrogen bonds with residues A48 and A49, which are known to be critical for ligand recognition.^16^ The warhead of **A1’** is positioned within a guanine-rich region where it forms multiple hydrogen bonds with G10, G32, G62 and G84. Given the established reactivity of phenylglyoxal with N1 and N2 of guanine at the Watson–Crick face, molecular docking results suggested that unpaired guanines G10, G11 and G62 adopt favorable conformations for covalent bond formation with the reactive warhead of **A1’** (Fig. S2C).

Subsequently, compound **A1** was synthesized and tested for its ability to modify covalently the FMN riboswitch. Matrix-assisted laser desorption/ionization-time of flight-based mass spectrometry (MALDI-TOF-MS) was used to characterize the covalent modification.^4^ Similar to a reported protocol,^14, 16^ the FMN riboswitch was constructed by annealing two synthesized strands, A and B (Fig. 2A). After reaction of **A1** with the riboswitch, the samples were denatured with 4 M urea to eliminate detection of non-covalent interactions followed by a clean-up and MS analysis (Fig. 2A). The denaturation conditions are critical to remove any binding compounds from detection as a covalent adduct in the MALDI-TOF-MS (Fig. S3).

**Figure 2.**
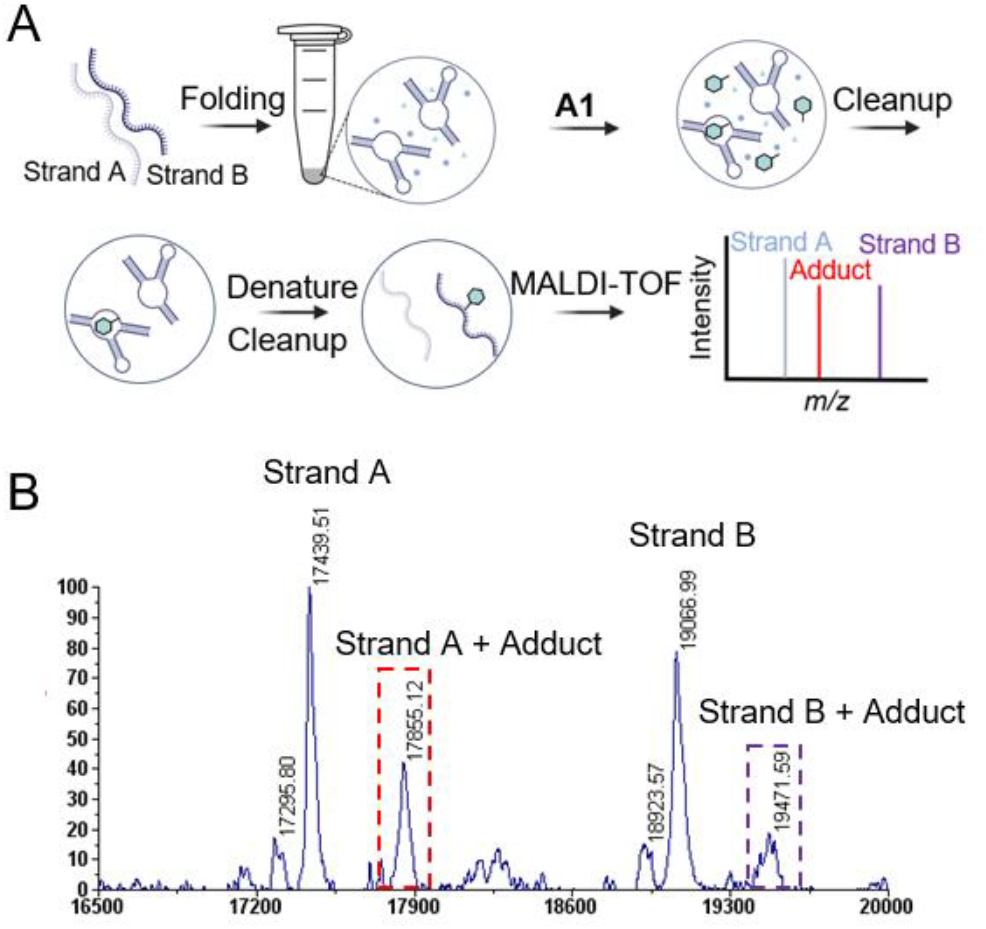
Mass spectrometric characterization of the covalent modification of the FMN riboswitch by A1. **A)** Schematic overview of the MALDI-TOF-based assay for evaluating covalent modification of FMN riboswitch by compound **A1. B)** A representative MALDI-TOF mass spectrum for the reaction of **A1** (200 μM) with FMN riboswitch (10 μM) at 37 °C for 12 h. Each peak is labeled with its corresponding mass-to-charge ratio (m/z). The peaks corresponding to the covalently modified strands A and B of the riboswitch are highlighted with red and purple dashed boxes, respectively. covalent small-molecule medicines that target RNA, as has been demonstrated in other systems.^3, 17^

The extent of covalent modification was quantified by integrating the peak areas corresponding to the modified and unmodified RNA. In addition to the unmodified strands, adducts were detected, corresponding to the mass expected for both strands A and B if the riboswitch construct was modified by **A1**. After 12 h, 200 µM of **A1** modified ∼33% and ∼24% of the strands A and B of the riboswitch (10 µM), respectively (Fig. 2B). In contrast, probe **P1**, which lacks a binding scaffold, modified ∼6% and ∼4% of the strands A and B under the same conditions (Fig. S4). Moreover, **A1** showed dose-dependent modification. As the concentration of **A1** increased from 40 μM to 200 μM, the extent of modification of strand A rose from 10 ±3% to 34±1% (Fig. S5). Similarly, a time-dependent increase in covalent modification was observed at a concentration of 100 μM, ranging for 15±3% at 3 h to 31±2% at 24 h (Fig. S6). These results demonstrated that **A1** preferentially and effectively modified strand A of the FMN riboswitch in a dose and time-dependent manner, in accordance with the structure guided design.

To support occupation of the FMN riboswitch binding pocket by **A1**, a competition assay was performed between **A1** and the non-covalent ligands **FMN** and riboflavin, the **FMN** precursor that lacks the phosphate and exhibits a moderate binding affinity (∼40 μM), ∼1000-fold weaker than that of **FMN** (Fig. S7A).^15^ Preincubation of **FMN** (50 μM, 15 min) completely abolished covalent modification of the FMN riboswitch (10 μM) by **A1** (100 μM, 8 h) (Fig. S7B, C). In line with its reduced binding affinity, preincubation of riboflavin at a concentration of 150 μM resulted in a significant decrease, but not complete ablation, of covalent modification induced by 100 µM **A1** (Fig. S7D). These results demonstrate that **FMN** and riboflavin compete with **A1** in the FMN binding pocket and limit access to and reaction with the unpaired guanines, supporting that the covalent modification is driven by induced proximity.

The guanine-derived imidazopurinone adduct formed by glyoxal derivatives has been reported to impede reverse transcriptase (RT), resulting in truncated complementary DNA (cDNA) strands.^9^ To identify the sites of reaction, an RT stop assay was carried out as described previously using SuperScript III enzyme and a strand A- or strand B-specific primer (Fig. 3A).^17^ The resulting cDNA products were analyzed by fragment analyzer capillary electrophoresis (CE) for quantitative determination of RT stop sites using customized ladders consisting of the target cDNA fragments. Compared with DMSO controls, treatment with compound **A1** (100 μM) and probe **P1** (1 mM) induced a distinct truncated cDNA product on strand A with a length of ∼43 nucleotides (nt), which mapped to the position of G11 on strand A of the FMN riboswitch (Fig. 3B). Comparable levels of the truncated cDNA were observed in **A1** and **P1**-treated samples, despite **A1** being present at a 10-fold lower concentration. No distinct truncated cDNA was detected on strand B upon treatment with either **A1** or **P1** (Fig. S8A).

**Figure 3.**
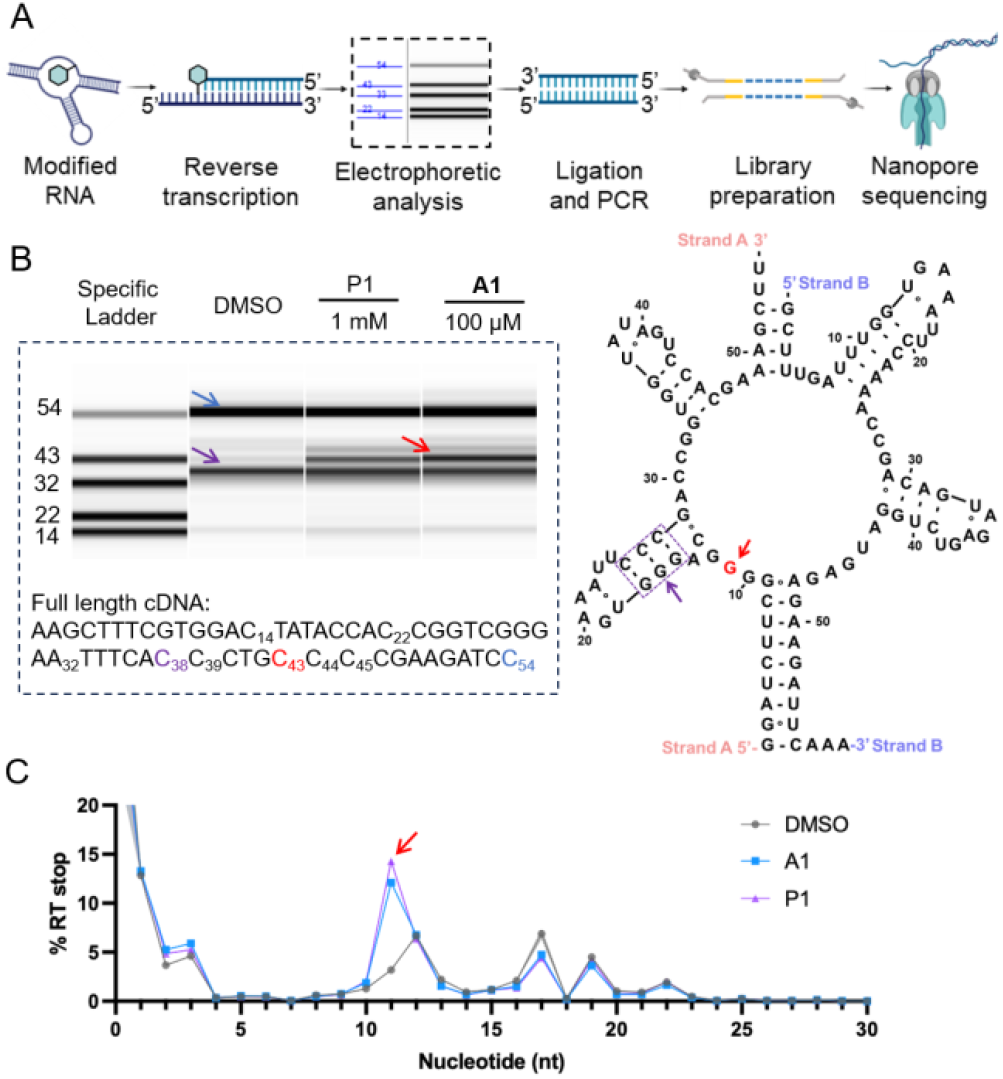
Identification of the sites of covalent FMN riboswitch modification by A1. **A)** Schematic overview of the detection of covalently modified RNA nucleotides, which induce reverse transcriptase (RT) stops, by nanopore sequencing. **B)** Left: Representative fragment analyzer traces for cDNA products from strand A of the constructed FMN riboswitch. The full length of cDNA product of strand A is 54 nt (blue arrow). Treatment with compound **A1** (100 μM) and **P1** (1 mM) for 8 h results in a distinct RT stop product of ∼43 nt (red arrow). An RT stop product at ∼38 nt is observed for all treatment groups, including DMSO (purple arrow). The length of RT stops was determined using a custom ladder (“Specific Ladder”) comprised of DNA sequences identical to the cDNA products of the corresponding size (Table S1). Right: RT stops mapped onto the secondary structure of FMN riboswitch (right). **C)** Representative RT stop profile of strand A reacted with **A1** (100 μM) and **P1** (1 mM) obtained from nanopore sequencing.

To further resolve the cross-linking sites of **A1**, truncated cDNA products were sequenced using nanopore sequencing technology (Fig. 3A).^17^ The results revealed that both **A1** and **P1** induced RT stops at nucleotide G11 of strand A (Fig. 3D), although **P1** required a 10-fold higher concentration to achieve a comparable percentage of RT stop. Interestingly, a weak RT stop was also detected at G6 of strand B (Fig. S8B, C), corresponding to G62 in the FMN riboswitch structure (Fig. 1B, S2C). This cDNA product was not detected in previous fragment analysis because its signal was obscured by the full-length cDNA.

These findings indicate that **A1** modifies the FMN riboswitch by reacting with G11 of strand A and G6 of strand B. The differing intensities of RT stops are consistent with the observed differences in covalent modification between the two strands. Furthermore, these results align with the prior docking analysis of **A1’**, which predicted that the covalent warhead could form hydrogen bonds and adopt a favorable reactive conformation at both G11 and G62 (G6 of strand B). Together, these data support a mechanism in which the compound specifically targets unpaired guanines near the FMN-binding site through proximity-induced covalent reactivity. This proximity induced reactivity strategy could enable the development of covalent small-molecule medicines that target RNA, as has been demonstrated in other systems.^3, 17^

Finally, to test the biological activity of **A1**, a LacZ reporter assay was performed in *Bacillus subtilis PY79*. The *B. subtilis ribD* leader sequence, which regulates gene expression at the transcriptional level, was fused to a LacZ reporter under an isopropylthio-*β*-galactoside (IPTG)-inducible Pspank promoter and integrated into the genome at the *amyE* locus (Fig. S9).^13, 18^ Modulation of the FMN riboswitch was assessed using the Miller assay, where o-nitrophenyl β-d-galactopyranoside (ONPG) cleavage yields a yellow product, o-nitrophenol (ONP), as a readout of riboswitch-controlled gene expression (Fig. 4A).^19^

**Figure 4.**
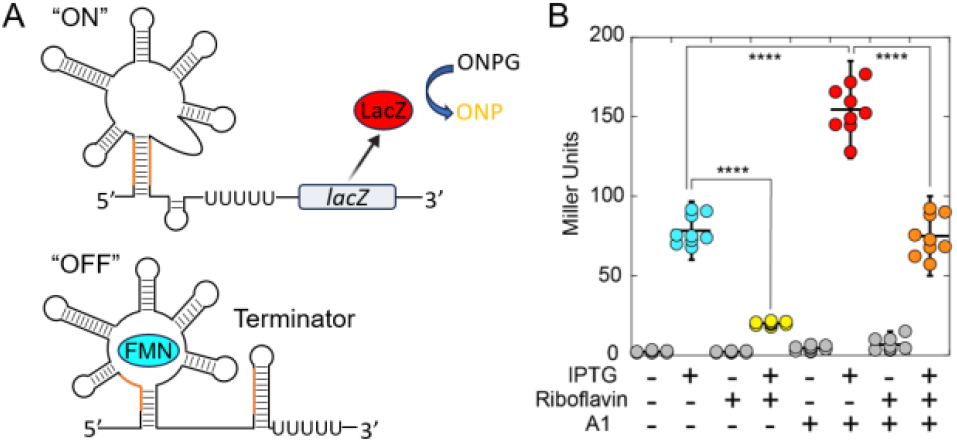
Biological activity of A1 by an FMN riboswitch reporter system in *Bacillus subtilis*. **A)** Simplified schematic of the RFN-mediated regulation of an mRNA bearing a *lacZ* reporter gene. Protein LacZ cleaves 2-nitrophenyl-β-D-galactopyranoside (ONPG) to afford ortho-nitrophenol (ONP; yellow). **B)** Miller units represent LacZ activity, reflecting relative gene expression levels upon reporter induction by IPTG (1 mM) in the presence or absence of riboflavin (100 µM; yellow) and **A1** (200 µM; red).

As expected, when expression of the reporter is induced by addition of IPTG but riboflavin is absent, a reproducibly higher level of LacZ expression was observed (cyan; Fig. 4B).^13^ Addition of riboflavin to the growth medium leads to its intracellular conversion to **FMN**, which switched the riboswitch conformation such that the terminator structure was formed (“OFF”, Fig. 4A), thereby reducing expression of the reporter (cyan and yellow; Fig. 4B).

Despite its structural similarity to riboflavin, which attenuates reporter expression, the presence of 200 µM of **A1** has a stimulatory effect on reporter expression (red; Fig. 4B). That is, **A1** acts as an agonist, rather than an antagonist as **FMN** does. The previous non-covalent docking predicts **A1’** could bind within the FMN binding pocket and assume a pose like **FMN** (Fig. S2B). This suggests that covalent modification of riboswitch by **A1** may promote and stabilize an alternative secondary structure (Fig 4A), resulting in an anti-terminated state. When both riboflavin (100 µM) and **A1** (200 µM) were added to the growth medium, a reduction of gene expression was observed (orange; Fig. 4B), similar to that observed upon induction of the reporter alone (cyan; Fig. 4B). These data suggest that these two compounds competitively bind to the riboswitch in cells, as was observed *in vitro* (Fig. S7).

Collectively, these findings highlight the ability of covalent small molecules to modulate RNA function by targeting specific residues within RNA structures. Notably, **A1** represents the first riboswitch-targeting compound that elicits the opposite effect of its natural effector molecule, underscoring that this approach can afford small molecules that modulate RNA function.

RNA-targeted covalent small molecules remain a largely underexplored area in drug discovery, due to the limited availability of electrophilic warheads that can react selectively with defined RNA nucleophiles. In this study, we demonstrate that phenylglyoxal can be incorporated into a rationally designed ligand to covalently modify unpaired guanines in structured RNA elements. Using the FMN riboswitch as a model system, we showed that the phenylglyoxal-based compound **A1** modifies specific guanine residues (G11 and G62) proximal to the FMN-binding site through a proximity-driven mechanism. This is guided by the ligand scaffold, which binds within the FMN pocket and orients the warhead toward nucleophilic residues.

Docking analysis, mass spectrometry, RT-stop mapping, and nanopore sequencing all support a model in which **A1** modifies the RNA *via* direct covalent bond formation at guanines exposed in the tertiary structure of the riboswitch. Importantly, only weak covalent reactivity is observed with the phenylglyoxal probe alone (**P1**), highlighting the requirement for scaffold-induced proximity to achieve selective RNA modification. The competition experiments with **FMN** and riboflavin further confirm that covalent labeling is driven by binding-site engagement rather than non-specific reactivity.

In addition to the biochemical characterization, compound **A1** was found to modulate gene expression in a cellular riboswitch reporter assay. Unlike **FMN**, which represses LacZ expression by stabilizing the terminated RNA structure, **A1** promotes LacZ expression, suggesting that it favors the anti-terminated conformation. The ability of **A1** to stimulate anti-termination effect underscores the potential for covalent RNA ligands to yield previously unknown mechanisms of functional modulation distinct from naturally occurring ligands or conventional small molecules.

Notably, **A1** is the first reported riboswitch-targeting compound that covalently modifies RNA and acts as a negative antagonist by eliciting the opposite effect of the natural effector molecule. These results establish a proof-of-concept for using structure-guided design to develop selective covalent ligands that modulate RNA function through induced reactivity. Unpaired guanines are widespread in regulatory and disease-associated RNAs, for examples, the thiamine pyrophosphate (TPP) riboswitch, which contains unpaired guanines near the ligandbinding site;^20^ expanded r(G_4_C_2_) and r(CGG) repeats featuring unpaired 1×1 nucleotide GG internal loops;^10, 21^ and the structured internal ribosomal entry site (IRES) of *MYC* mRNA and the iron-responsive element (IRE) of α-synuclein (*SNCA*) mRNA, both of which contain multiple unpaired guanines.^22^ Therefore, this approach offers a broadly applicable framework for RNA-targeted drug discovery. Follow-up studies will include demonstrating more general applicability of this reactive module and understanding the structure and mechanistic underpinnings of the mode of action. This study, however, demonstrates that structure-guided design can be used to imbue RNA-binding small molecules with specific covalent reactivity.

## ASSOCIATED CONTENT

### Supporting Information

The following data can be found in the Supporting Information. (i) Figures S1™S9; (ii) materials and methods; and (iii) synthetic methods and characterization. The Supporting Information is available free of charge on the ACS Publications website.

## AUTHOR INFORMATION

### Corresponding Author

**Matthew D. Disney** − Department of Chemistry, The Scripps Research Institute; Department of Chemistry, The Herbert Wertheim UF Scripps Institute for Biomedical Innovation & Technology, Jupiter, Florida 33458, United States., **Robert T. Batey** − Department of Molecular, Cellular and Developmental Biology; Department of Biochemistry, University of Colorado, Boulder, Colorado 80309-0596, United States.

### Authors

**Chungen Li** –*Department of Chemistry, The Herbert Wertheim UF Scripps Institute for Biomedical Innovation & Technology, Jupiter, FL 33458, United States*.

**Xueyi Yang** –*Department of Chemistry, The Scripps Research Institute, Jupiter, Florida 33458, United States*.

**Kyle A. Dickerson** *– Department of Molecular, Cellular and Developmental Biology, University of Colorado, Boulder, Colorado 80309-0596, United States*.

**Patrick R. A. Zanon** – *Department of Chemistry, The Herbert Wertheim UF Scripps Institute for Biomedical Innovation & Technology, Jupiter, FL 33458, United States*.

**Noah A. Springer** – *Department of Chemistry, The Scripps Research Institute, Jupiter, Florida 33458, United States*.

**Jielei Wang** – *Department of Chemistry, The Scripps Research Institute, Jupiter, Florida 33458, United States*.

**Yilin Jia** – *Department of Chemistry, The Scripps Research Institute, Jupiter, Florida 33458, United States*.

**Nikhil C. Munshi –** *Department of Medical Oncology, Jerome Lipper Multiple Myeloma Center, Dana-Farber Cancer Institute, Boston, MA 02215, United States*

### Author Contributions

The manuscript was written through contributions of all authors. All authors have given approval to the final version of the manuscript.

### Notes

The authors declare no competing financial interest.

## ACKNOWLEDGMENT

This work was supported by the U.S. National Institutes of Health (R35 NS116846 (to M.D.D.), R01 P0377752 (to M.D.D. and N.C.M.) and R35 GM152029 (to R.T.B.).

## ABBREVIATIONS

cDNA: complementary DNA;
CE: capillary electrophoresis;
FMN: flavin mononucleotide;
IPTG: isopropylthio-*β*-galactoside;
IRE: iron-responsive element;
IRES: internal ribosomal entry site;
MALDI-TOF-MS: matrix-assisted laser desorption/ionization-time of flight-based mass spectrometry; m/z mass-to-charge ratio;
NPG: o-nitrophenol;
ONPG: o-nitrophenyl β-d-galactopyranoside;
RT: reverse transcriptase;
TAMRA: tetramethylrhodamine;
TPP: thiamine pyrophosphate.

## TOC Graphic

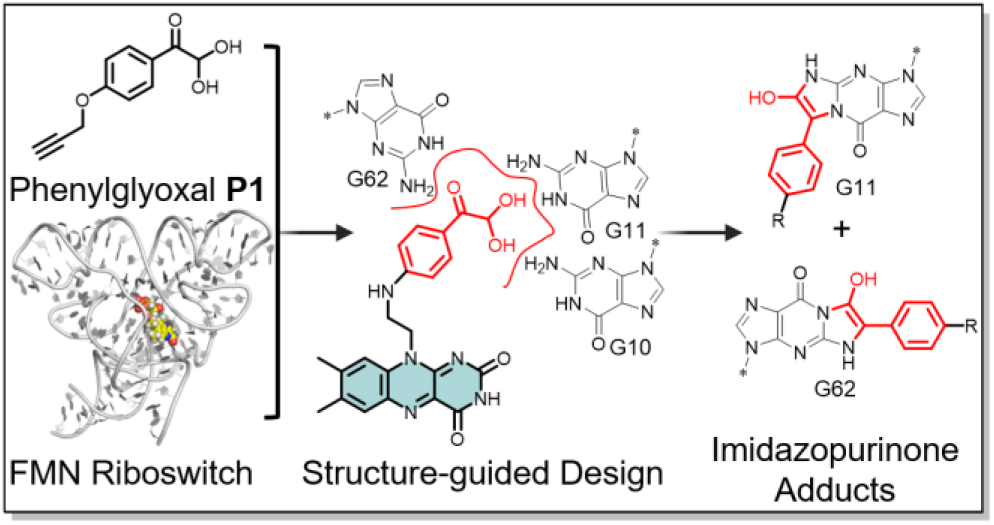

## REFERENCES

(1) Raouf, Y. S. Covalent Inhibitors: To Infinity and Beyond. J Med Chem 2024, 67 (13), 10513–10516. DOI: 10.1021/acs.jmedchem.4c01308. Boike, L.; Henning, N. J.; Nomura, D. K. Advances in covalent drug discovery. Nat Rev Drug Discov 2022, 21 (12), 881–898. DOI: 10.1038/s41573-022-00542-z From NLM Medline. Disney, M. D. Targeting RNA with Small Molecules To Capture Opportunities at the Intersection of Chemistry, Biology, and Medicine. J Am Chem Soc 2019, 141 (17), 6776–6790. DOI: 10.1021/jacs.8b13419.

(2) Guan, L.; Disney, M. D. Covalent small-molecule-RNA complex formation enables cellular profiling of small-molecule-RNA interactions. Angew Chem Int Ed Engl 2013, 52 (38), 10010–10013. DOI: 10.1002/anie.201301639. Yang, W. Y.; Wilson, H. D.; Velagapudi, S. P.; Disney, M. D. Inhibition of Non-ATG Translational Events in Cells via Covalent Small Molecules Targeting RNA. J Am Chem Soc 2015, 137 (16), 5336–5345. DOI: 10.1021/ja507448y.

(3) Rzuczek, S. G.; Colgan, L. A.; Nakai, Y.; Cameron, M. D.; Furling, D.; Yasuda, R.; Disney, M. D. Precise small-molecule recognition of a toxic CUG RNA repeat expansion. Nat Chem Biol 2017, 13 (2), 188–193. DOI: 10.1038/nchembio.2251.

(4) Springer, N. A.; Zanon, P. R. A.; Taghavi, A.; Sung, K.; Disney, M. D. Discovery of RNA-reactive small molecules guides design of electrophilic modules for RNA-specific covalent binders. bioRxiv 2025. DOI: 10.1101/2025.04.22.649986.

(5) McDonald, R. I.; Guilinger, J. P.; Mukherji, S.; Curtis, E. A.; Lee, W. I.; Liu, D. R. Electrophilic activity-based RNA probes reveal a self-alkylating RNA for RNA labeling. Nat Chem Biol 2014, 10 (12), 1049–1054. DOI: 10.1038/nchembio.1655. Bereiter, R.; Flemmich, L.; Nykiel, K.; Heel, S.; Geley, S.; Hanisch, M.; Eichler, C.; Breuker, K.; Lusser, A.; Micura, R. Engineering covalent small molecule-RNA complexes in living cells. Nat Chem Biol 2025, 21 (6), 843–854. DOI: 10.1038/s41589-024-01801-3.

(6) Balaratnam, S.; Rhodes, C.; Bume, D. D.; Connelly, C.; Lai, C. C.; Kelley, J. A.; Yazdani, K.; Homan, P. J.; Incarnato, D.; Numata, T.; et al. A chemical probe based on the PreQ1 metabolite enables transcriptome-wide mapping of binding sites. Nat Commun 2021, 12 (1), 5856. DOI: 10.1038/s41467-021-25973-x.

(7) Osborne., M. R.; Wilman., D. E. V.; Lawley., P. D. Alkylation of DNA by the nitrogen mustard bis(2-chloroethyl)methylamine. Chem Res Toxicol 1995, 8 (2), 316–320. DOI: 10.1021/tx00044a018. Yu, H.; Hiroki, U.; Gakushi, K.; Shingo, N.; Masaaki, K.; Hideyuki, S. A reaction mechanism-based prediction of mutagenicity: α-halo carbonyl compounds adduct with DNA by S_N_2 reaction. J Toxicol Sci 2018, 43 (3), 203–211. DOI: 10.2131/jts.43.203.

(8) Robert, S.; Cohen., B. I.; Shiuey., S.-J.; Hans, M. On the reaction of guanine with glyoxal, pyruvaldehyde, and kethoxal, and the structure of the acylguanines. A new synthesis of N^2^-Alkylguanines. Biochemistry 1969, 8, 238–245. DOI: 10.1021/bi00829a034. Masaaki, K.; Sohsuke, K.; Kazue, S. A chemiluminescence derivatization method for detecting nucleic acids and DNA probes using a trimethoxyphenylglyoxal reagent that recognizes guanine. Analytica Chimica Acta 1999, 381, 155–163.

(9) Mitchell, D.; Ritchey, L. E.; Park, H.; Babitzke, P.; Assmann, S. M.; Bevilacqua, P. C. Glyoxals as in vivo RNA structural probes of guanine base-pairing. RNA 2018, 24 (1), 114–124. DOI: 10.1261/rna.064014.117.

(10) Wang, Z. F.; Ursu, A.; Childs-Disney, J. L.; Guertler, R.; Yang, W. Y.; Bernat, V.; Rzuczek, S. G.; Fuerst, R.; Zhang, Y. J.; Gendron, T. F.; et al. The hairpin form of r(G_4_C_2_)^exp^ in c9ALS/FTD is repeat-associated non-ATG translated and a target for bioactive small molecules. Cell Chem Biol 2019, 26 (2), 179–190 e112. DOI: 10.1016/j.chembiol.2018.10.018.

(11) Kolb, H. C.; Finn, M. G.; Sharpless, K. B. Click chemistry: Diverse chemical function from a few good reactions. Angew Chem Int Ed Engl 2001, 40 (11), 2004–2021. Kolb, H. C.; Sharpless, K. B. The growing impact of click chemistry on drug discovery. Drug Discov Today 2003, 8 (24), 1128–1137.

(12) Kavita, K.; Breaker, R. R. Discovering riboswitches: the past and the future. Trends Biochem Sci 2023, 48 (2), 119–141. DOI: 10.1016/j.tibs.2022.08.009.

(13) Winkler, W. C.; Cohen-Chalamish, S.; Breaker, R. R. An mRNA structure that controls gene expression by binding FMN. Proc Natl Acad Sci U S A 2002, 99 (25), 15908–15913. DOI: 10.1073/pnas.212628899.

(14) Vicens, Q.; Mondragon, E.; Batey, R. T. Molecular sensing by the aptamer domain of the FMN riboswitch: a general model for ligand binding by conformational selection. Nucleic Acids Res 2011, 39 (19), 8586–8598. DOI: 10.1093/nar/gkr565.

(15) Serganov, A.; Huang, L.; Patel, D. J. Coenzyme recognition and gene regulation by a flavin mononucleotide riboswitch. Nature 2009, 458 (7235), 233–237. DOI: 10.1038/nature07642.

(16) Vicens, Q.; Mondragon, E.; Reyes, F. E.; Coish, P.; Aristoff, P.; Berman, J.; Kaur, H.; Kells, K. W.; Wickens, P.; Wilson, J.; et al. Structure-activity relationship of flavin analogues that target the flavin mononucleotide riboswitch. ACS Chem Biol 2018, 13 (10), 2908–2919. DOI: 10.1021/acschembio.8b00533.

(17) Yang, X.; Wang, J.; Springer, N. A.; Zanon, P. R.; Jia, Y.; Su, X.; Disney, M. D. Mapping small molecule–RNA binding sites via Chem-CLIP synergized with capillary electrophoresis and nanopore sequencing. Nucleic Acids Research 2025, 53 (6), gkaf231.

(18) Wickiser, J. K.; Winkler, W. C.; Breaker, R. R.; Crothers, D. M. The speed of RNA transcription and metabolite binding kinetics operate an FMN riboswitch. Mol Cell 2005, 18 (1), 49–60. DOI: 10.1016/j.molcel.2005.02.032.

(19) Miller, J. H. Experiments in Molecular Genetics; Cold Spring Harbor Laboratory, 1972.

(20) Zeller, M. J.; Favorov, O.; Li, K.; Nuthanakanti, A.; Hussein, D.; Michaud, A.; Lafontaine, D. A.; Busan, S.; Serganov, A.; Aube, J.; et al. SHAPE-enabled fragment-based ligand discovery for RNA. Proc Natl Acad Sci U S A 2022, 119 (20), e2122660119. DOI: 10.1073/pnas.2122660119.

(21) Disney, M. D.; Liu, B.; Yang, W. Y.; Sellier, C.; Tran, T.; Charlet-Berguerand, N.; Childs-Disney, J. L. A small molecule that targets r(CGG)(exp) and improves defects in fragile X-associated tremor ataxia syndrome. ACS Chem Biol 2012, 7 (10), 1711–1718. DOI: 10.1021/cb300135h.

(22) Zhang, P.; Park, H. J.; Zhang, J.; Junn, E.; Andrews, R. J.; Velagapudi, S. P.; Abegg, D.; Vishnu, K.; Costales, M. G.; Childs-Disney, J. L.; et al. Translation of the intrinsically disordered protein alpha-synuclein is inhibited by a small molecule targeting its structured mRNA. Proc Natl Acad Sci U S A 2020, 117 (3), 1457–1467. DOI: 10.1073/pnas.1905057117. Tong, Y.; Lee, Y.; Liu, X.; Childs-Disney, J. L.; Suresh, B. M.; Benhamou, R. I.; Yang, C.; Li, W.; Costales, M. G.; Haniff, H. S.; et al. Programming inactive RNA-binding small molecules into bioactive degraders. Nature 2023, 618 (7963), 169–179. DOI: 10.1038/s41586-023-06091-8.

