## Supporting Information for "Structure-Guided Design of a Bioactive Covalent Small Molecule Targeting a Riboswitch"

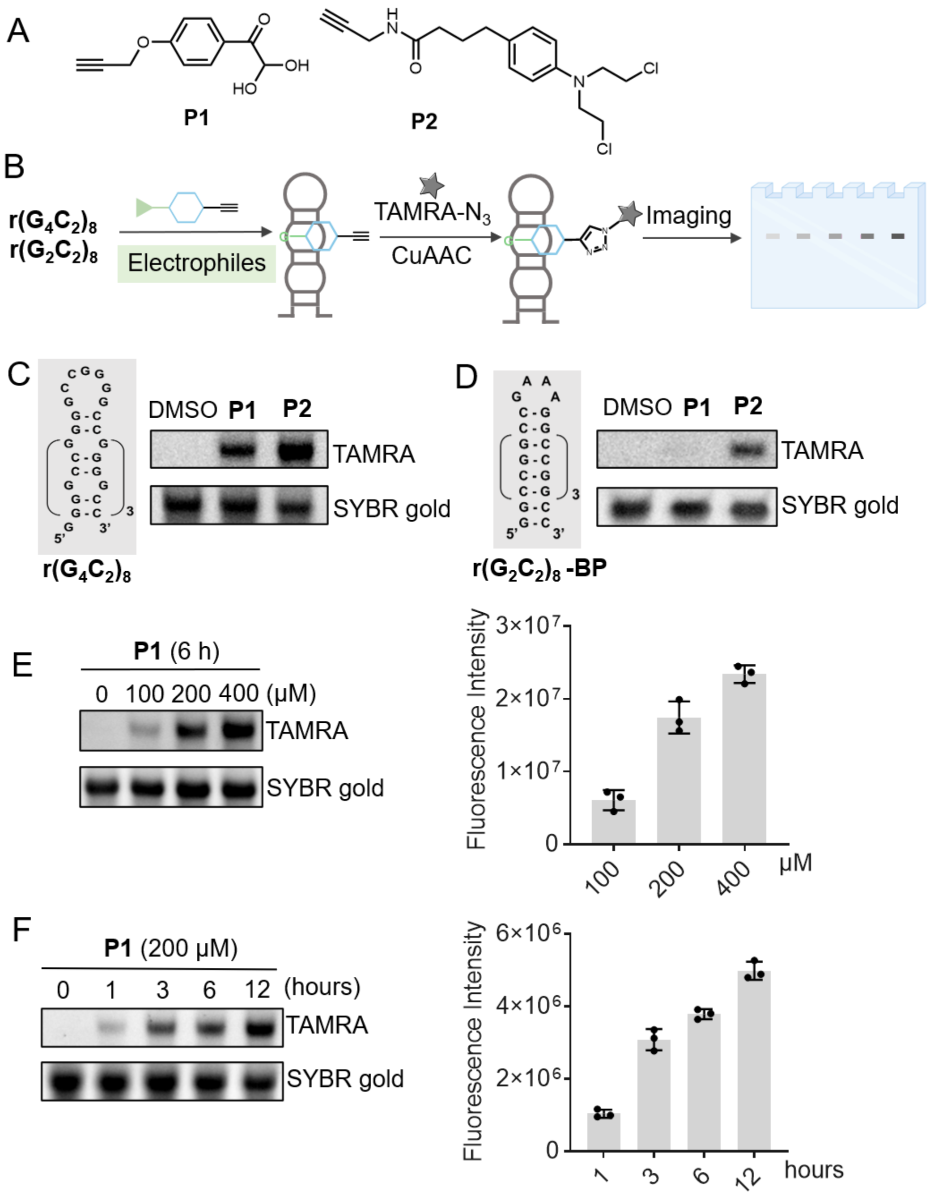


**Figure S1**. **Characterization of phenylglyoxal probe targeting unpaired guanines in structured RNAs.** **A)** The chemical structures of phenylglyoxal **P1** and chlorambucil **P2**. **B)** Schematic representation of the TAMRA labeling assay workflow. **C)** Secondary structure of r($G_4C_2$)$_8$ (left) and TAMRA labeling of r($G_4C_2$)$_8$ by **P1** (200 μM) and **P2** (200 μM) at 37 °C for 6 h (right). **D)** Secondary structure of base paired r($G_2C_2$)$_8$ (left) and TAMRA labeling by **P1** and **P2** (200 μM) at 37 °C for 6 h (right). **E)** Dose-dependent labeling of r($G_4C_2$)$_8$ by **P1** (100, 200 and 400 μM) at 37 °C for 6 h (left), with quantification of fluorescence intensity from three independent experiments (right). **F)** Time-dependent labeling of r($G_4C_2$)$_8$ by **P1** (200 μM) at 37 °C (left) and corresponding quantification of fluorescence intensity (n = 3, right).

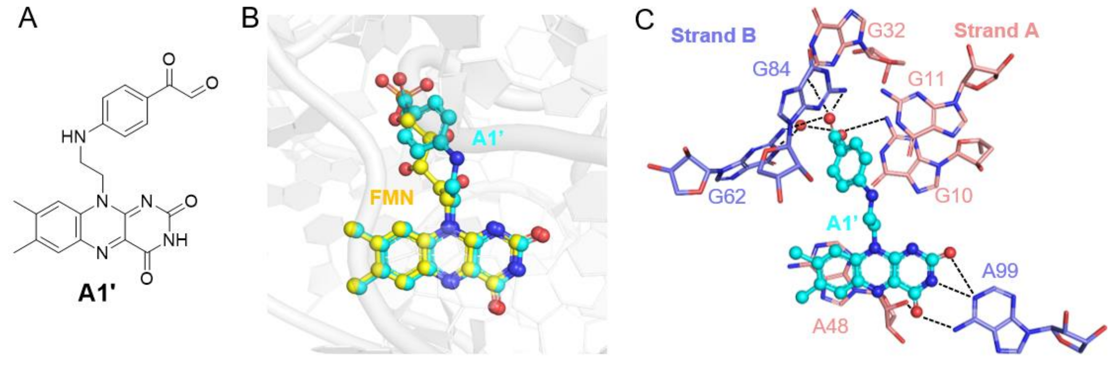


**Figure S2. Molecular docking analysis of A1′ within the FMN riboswitch binding pocket.** **A)** Chemical structure of **A1′**, the dehydration form of **A1**. **B)** Alignment of **FMN** (yellow stick) with **A1’** (cyan stick) within the FMN-binding pocket. **C)** Predicted binding mode of **A1’** (cyan stick) within the FMN riboswitch. Hydrogen bonds are indicated by black dashed lines.

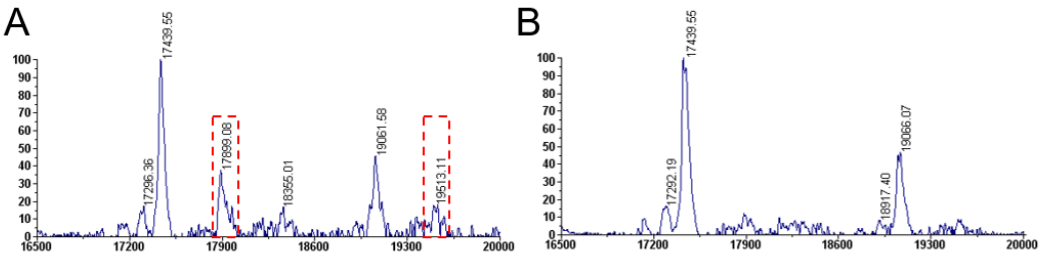


**Figure S3. Denaturation conditions eliminate detection of non-covalent interactions in MALDI-TOF MS**. Representative MALDI-TOF MS spectra of FMN riboswitch treated with non-covalent **FMN** (200 μM, 12 h) under native (**A**) and denaturing (**B**) conditions. Denaturation was performed with 4 M urea at 37 °C for 15 min. The peaks corresponding to non-covalent interactions by **FMN** are highlighted with red dashed box.

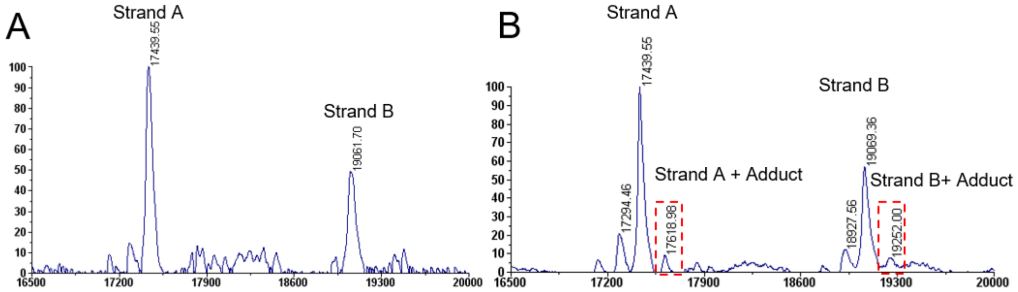


**Figure S4**. Representative MALDI-TOF mass spectra of the FMN riboswitch (10 μM) treated with DMSO (**A**) or **P1** (200 μM) (**B**) at 37 °C for 12 h. The peaks corresponding to covalently modified strands are highlighted with red dashed box.

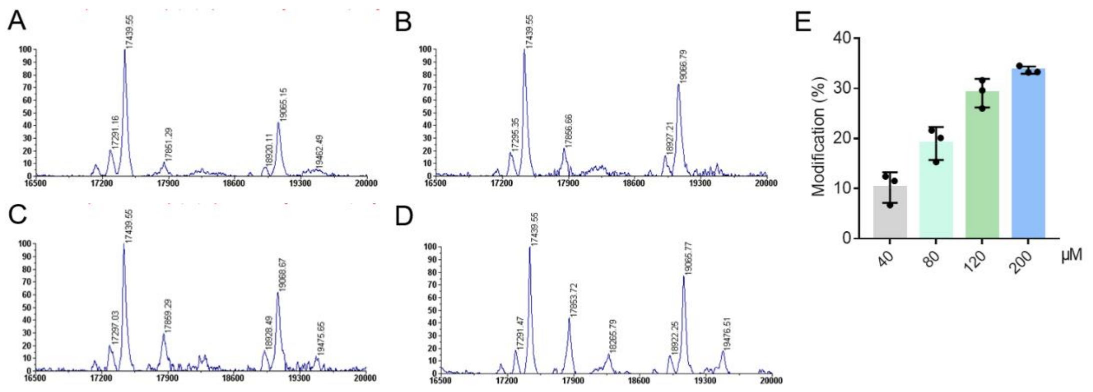


**Figure S5. Dose-dependent covalent modification of the FMN riboswitch by A1.** (**A–D**) Representative MALDI-TOF mass spectra of FMN riboswitch (10 μM) treated by **A1** at concentrations of 40 μM (**A**), 80 μM (**B**), 120 μM (**C**), and 200 μM (**D**) at 37 °C for 12 h. **E)** Quantification of covalent modification based on the integrated peak areas of unmodified and modified strand A (n = 3).

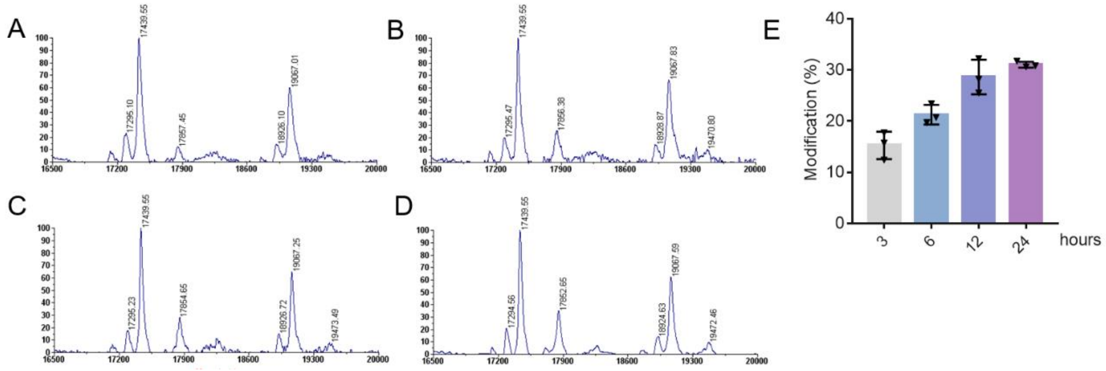


**Figure S6. Time-dependent covalent modification of the FMN riboswitch by A1.** (**A–D**) Representative MALDI-TOF mass spectra of FMN riboswitch (10 μM) treated by **A1** (100 μM) at 37 °C for 3 h (**A**), 6 h (**B**), 12 h (**C**), and 24 h (**D**). **E)** Quantification of covalent modification based on the integrated peak areas of unmodified and modified strand A (n = 3).

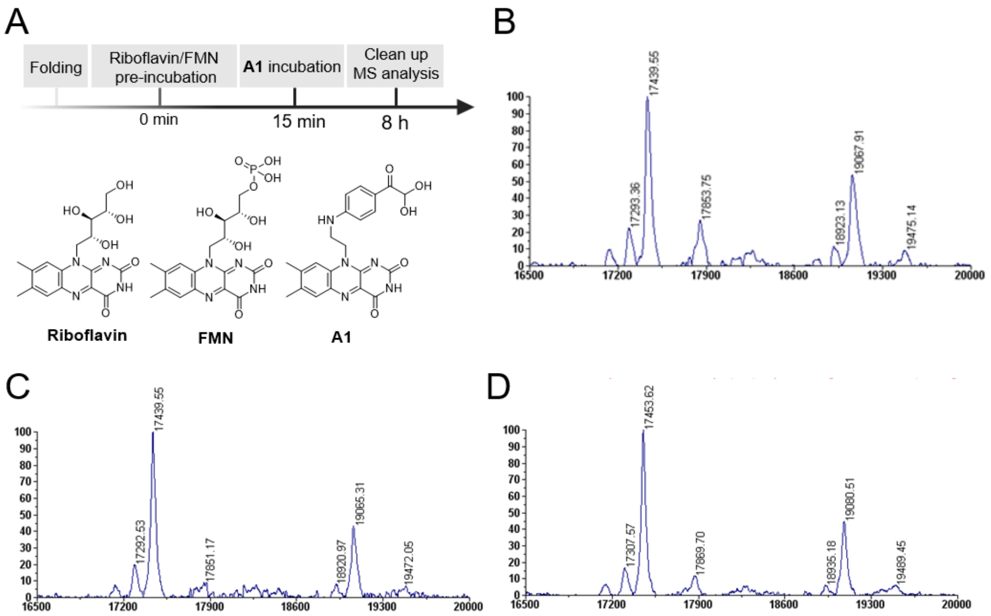


**Figure S7. Competitive inhibition of A1-induced modification by FMN and riboflavin.** **A**) Schematic overview of the competition assay. **B–D**) Representative MALDI-TOF mass spectra of the FMN riboswitch (10 μM) pre-incubated with DMSO (**B**), **FMN** (50 μM, **C**), or **riboflavin** (150 μM, **D**) at 37 °C for 15 min, followed by treatment with **A1** (100 μM) for another 8 h.

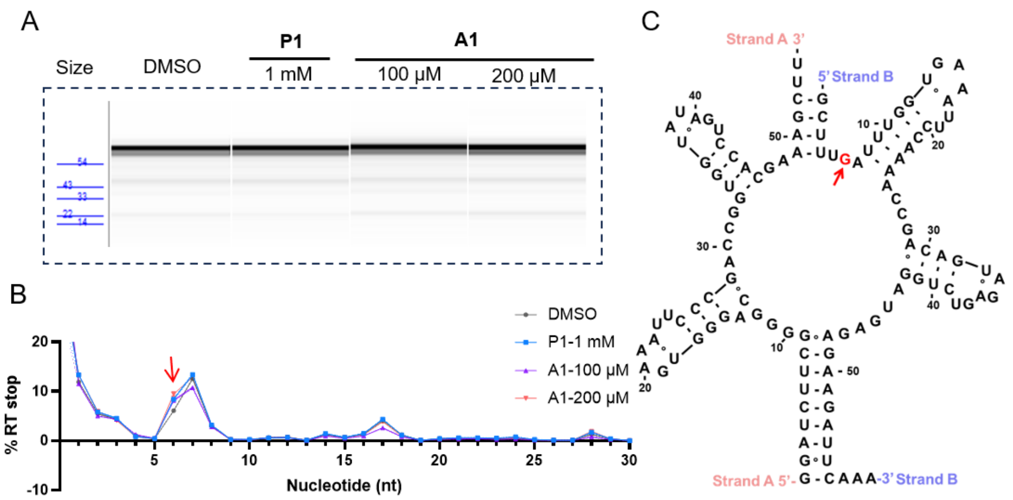


**Figure S8. Identification of the A1 crosslinking site on FMN riboswitch strand B. A)** Representative fragment analyzer traces for cDNA products from strand B treated with compound **A1** (100 μM, 200 μM) and **P1** (1 mM) for 8 h. **B)** Representative RT stop profile of strand B reacted with **A1** (100 μM, 200 μM) and **P1** (1 mM) obtained from nanopore sequencing (n = 3). **C)** RT stops mapped onto the secondary structure of FMN riboswitch, with the RT stop site G6 highlighted in red.

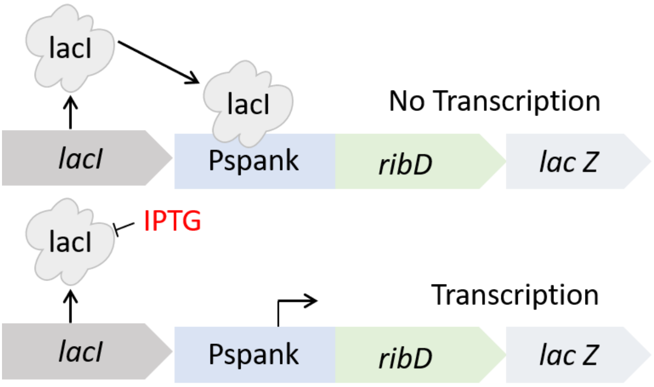


**Figure S9.** Simplified schematic of the reporter system for the bioactivity assay.

**Table S1.** Sequences of oligonucleotides used in this study.

| Name | Sequence [α] | Assay |
|---|---|---|
| r($G_4C_2$)$_8$ RNA | 5'-GGGGCCGGGGCCGGGGCCGGGGCCGGGGCCG GGGCCGGGGCCGGGGCC-3' | TAMRA |
| r($G_2C_2$)$_8$ RNA | 5'-GGCCGGCCGGCCGGCCGGCCGGCCGGCCGGC C-3' | TAMRA |
| Riboswitch strand A | 5'-GGAUCUUCGGGGCAGGGUGAAAUUCCCGACCG GUGGUAUAGUCCACGAAAGCUU-3' | MALDI-TOF |
| Riboswitch strand B | 5'-GCUUUGAUUUGGUGAAAUUCCAAAACCGACAGU AGAGUCUGGAUGAGAGAAGAUUCAAA-3' | MALDI-TOF |
| Riboswitch strand A RT primer | 5'-AAGCTTTCGTGGAC-3' | RT-STOP |
| Riboswitch strand B RT primer | 5'-GAATCTTCTCTCATCCAGACTCTACTG-3' | RT-STOP |
| ssDNA ladder | 5'-AAGCTTTCGTGGACTATACCAC-3'<br>5'-AAGCTTTCGTGGACTATACCACCGGTCGGGAAT-3'<br>5'-AAGCTTTCGTGGACTATACCACCGGTCGGGAATTT CACCCTGC-3'<br>5'-AAGCTTTCGTGGACTATACCACCGGTCGGGAATTT CACCCTGCCCCGAAGATCC-3' | Analyzing RT cDNA products |
| Ligation adaptor | 5'-/5phos/AGATCGGAAGAGCGTCGTGTAG/3spc/-3' | Ligation before PCR |
| PCR primer of strand A | Forward: 5'-CTACACGACGCTCTTCCGATCT-3'<br>Reverse: 5'-AAGCTTTCGTGGACTATACCAC-3' | PCR |
| PCR primer of strand B | Forward: 5'-CTACACGACGCTCTTCCGATCT-3'<br>Reverse: 5'-GAATCTTCTCTCATCCAGACTCTACTG-3' | PCR |

[α] All RNA constructs were custom ordered from Horizon Discovery, and DNA constructs were ordered from Integrated DNA Technologies, Inc. (IDT).

### MATERIALS AND METHODS

#### *RNA folding*

*r($G_4C_2$)$_8$ and r($G_2C_2$)$_8$*

Chemically synthesized r($G_4C_2$)$_8$ and r($G_2C_2$)$_8$ were purchased from Horizon Discovery. The RNAs were dissolved in RNase-free water to a stock concentration of 100 μM. For folding, the RNA was diluted to 10 μM in 1× Folding Buffer (10 mM sodium phosphate, pH 7.0 and 100 mM LiCl). The RNAs solution was heated at 95 °C for 3 min and then immediately snap cooled on ice for at least 10 min. Folded RNAs were used immediately for the TAMRA labeling assay.

*FMN riboswitch*

The FMN riboswitch was constructed by two chemically synthesized strands A and B. Strand A (10 μM) and Strand B (10 μM) were mixed and folded in buffer containing 15 mM $MgCl_2$, 100 mM KCl, and 50 mM *N*-2-hydroxyethylpiperazine-*N*-2-ethane sulfonic acid (HEPES), pH 8.0. The mixture was heated at 95 °C for 3 min and cooled down to 25 °C by 3 °C/min. A refold was performed at 37 °C for 30 min prior to use.

#### *TAMRA labeling assay*

The TAMRA labeling assay was performed using a modified version of a previously described protocol.[1] Folded RNAs (10 μM) samples were incubated with probe compounds at 37 °C at the indicated concentrations and times to allow covalent modification. The modified RNA was cleaned up using RNAClean XP beads (Beckman Coulter, catalog # A63881) and eluted in RNase-free water (10 μL). The cleaned up RNA was then incubated with 5 μL of a “click mix” containing copper sulfate ($CuSO_4$, 20 mM, 0.1 μL), tris(3-hydroxypropyltriazolylmethyl)amine (THPTA, 20 mM, 0.5 μL), sodium ascorbate (100 mM, 0.4 μL), TAMRA-$N_3$ (20 mM, 0.1 μL) and RNase-free water (3.9 μL) at 37 °C for 1 h. The RNA was again cleaned up using RNAClean XP beads and eluted in RNase-free water. The TAMRA-labeled RNA was analyzed by a urea denaturing 11% (w/v) acrylamide gel, which was imaged by using a ChemiDoc™ MP System (excitation: 550 nm; emission: 565 nm). The gel was then post-stained with SYBR Gold (Invitrogen, catalog # S11494, 1:10000 dilution in water) for 10 min with gentle shaking at room temperature. SYBR Gold signal was then measured (excitation: 496 nm; emission: 539 nm). Quantification was performed by Image Lab.

#### MALDI-TOF-MS sample preparation

The folded FMN riboswitch RNA (10 μM) was incubated with compounds at 37 °C to allow covalent modification. The modified RNA was cleaned up using RNAClean XP beads and eluted in RNase-free water. To remove the tightly bound non-covalent compounds, the eluted RNA was denatured in 4 M urea solution at 37°C for 15 min. The mixture was cleaned up using RNAClean XP beads (Beckman Coulter, catalog # A63881), and the RNA was eluted in RNase-free water.

**MALDI-TOF-MS analysis**

Covalent modification of RNA by small molecules was characterized using MALDI-TOF mass spectrometry as previously described.[2] In brief, the cleaned up RNA (1 μL) was spotted onto a MALDI plate and air-dried at room temperature. Subsequently, 2′,4′,6′-Trihydroxyacetophenone monohydrate (THAP) matrix (1 μL, prepared as 50 mg/mL diammonium citrate and 18 mg/mL THAP in 1/1 water/acetonitrile) was then added to the dried spot and allowed to crystallize at room temperature. Mass spectra were acquired by an Applied Biosystems 4800 plus MALDI-TOF mass spectrometer (laser intensity: 7000-7400), accumulating sub-spectra with a signal-to-noise ratio threshold of >15. Data processing and spectral analysis were carried out using Applied Biosystems/SciEX Data Explorer software. The extent of covalent modification was quantified by integrating the peak areas corresponding to the modified and unmodified RNA. Modification percentage was calculated using the following equation: Modification (%) = [Adduct peak area / (Adduct peak area + Unmodified peak area)] × 100%.

**Docking**

Molecular docking was performed using Glide in extra precision (XP) mode.[3] The structure of **A1'** was prepared using LigPrep in Maestro.[3] The structure of FMN riboswitch (PDB: 2YIE) was prepared using the Protein Preparation Wizard. The receptor grid was generated centered on the FMN-binding pocket. Docking was performed with flexible ligand sampling. The resulting pose were visualized using the PyMOL Molecular Graphics System (Version 3.0 Schrödinger, LLC.).

**Reverse transcription of RNA**

The reverse transcription of covalently modified RNA was carried out by using SuperScript III (Invitrogen, catalog # 18-080-093) enzyme. To 500-800 ng RNA in RNase-free water (10.4 μL) was added 1 μL of 5 μM RT Primer of strands A or B of constructed FMN riboswitch (Table S1), 1 μL of borate buffer (500 mM potassium borate, pH 7.0) in a total volume of 12.4 μL. The mixture was heated at 60 °C for 5 min and then slowly cooled to 25 °C with a rate of 0.1°C/s in a thermocycler. After primer annealing, 1 μL of 10 mM dNTPs, 4 μL of 5×First Strand Buffer, 1 μL of 0.1 M dithiothreitol (DTT), 0.6 μL of DMSO, 0.5 μL of RNaseOUT (Invitrogen, catalog # 10777019), and 0.5 μL of SuperScript III were added in a total volume of 20 μL. The mixture was heated at 50 °C for 10 min, and then 85 °C for 5 min. Subsequently, the RT sample was cooled to room temperature, and 1 μL of RNase T1 (5 units) and 1 μL of RNase H (5 units) were added to digest the RNA. Following incubation at 37 °C for 30 min, the cDNA was purified using Ampure XP beads (Beckman Coulter, catalog # A63881) and eluted by Nanopure water.

**Analyzing reverse transcription (RT) cDNA products using a fragment analyzer**

cDNA products from RT were analyzed on an Agilent 5300 Fragment Analyzer following the manufacturer's protocol for the Small RNA Kit (Agilent, catalog # DNF-470-0275). The cDNA was denatured at 70 °C for 10 min and mixed with 18 μL of Small RNA Diluent (provided with the kit). Separation was performed with a 30 s pre-run at 11.5 kV, 50 s injection at 8.0 kV, and 45 min separation at 11.5 kV. A custom DNA ladder, composed of oligonucleotides (14–54 nt) with sequences identical to strand A of the riboswitch, was used to calibrate fragment sizes.

**Library preparation and nanopore sequencing**

For PCR amplification, cDNA generated by reverse transcription was ligated to an adaptor with the sequence of 5'-/5phos/AGATCGGAAGAGCGTCGTGTAG/3spc/-3'. The ligation was performed using CircLigase ssDNA Ligase (Biosearch Technologies, Catalog # CL4111K) according to the manufacturer's protocol. The ligated cDNA was then amplified by PCR using Phusion High-Fidelity DNA Polymerase (NEB, Catalog # M0530S), following the manufacturer's protocol. Sequencing libraries were prepared using the Native Barcoding Kit 96 V14 (Oxford Nanopore, catalog # SQK-NBD114.96) following the manufacturer's protocol. The library was sequenced using a MinION R10.4.1 flow cell (Oxford Nanopore, catalog # FLO-MIN114).

**RNA sequencing data analysis**

Nanopore sequencing data were base-called and demultiplexed using Dorado's 'basecaller' and 'demux' subcommands, respectively.[4] The base-called reads were first converted to FASTQ format using 'samtools fastq' and then aligned to the target sequence using 'bowtie2' with the '--local' option. The resulting SAM files were converted to BAM files and sorted using samtools.[5] A customed script was used to extract the alignment start site of each read on FMN strand A or strand B from each BAM file. The alignment starts sites minus one position, representing RT drop-off sites, were aggregated and plotted in Prism.

***Bacillus subtilis* reporter assay**

An FMN riboswitch reporter system was constructed in *Bacillus subtilis* strain PY79 to assess the bioactivity of compound A1 in live cells. The *B. subtilis ribD* riboswitch sequence was amplified from PY79 genomic DNA using the primers 5'-GCCGCAAGCTTAAGGACAAATGAATAAAGATTGTATCCTTC (underlined sequence is the riboswitch) and GAATCCGTAATCATGGTCATTTGTTTCCCTCCCCTCTTTT using a PCR reaction with Q5 polymerase (New England Biolabs). The *B. subtilis ribD* riboswitch DNA sequence (range 4053877 to 4054173, GenBank reference sequence ID CP026038.1) was fused to a downstream lacZ gene using a second round of PCR and the DNA inserted between the HindIII and SphI sites of pDR110. This places the mRNA containing the *ribD* riboswitch and *lacZ* gene under control of the IPTG-inducible Pspank promoter. Whole-plasmid Nanopore sequencing (Quintara Biosciences) was used to sequence-verify the plasmid. The sequence-verified plasmid was transformed into *B. subtilis* PY79 using natural competence and selection with spectinomycin.[6] Genomic integration at the *amyE* locus and sequence validation was achieved by Illumina whole genome sequencing at a minimum of 200Mbp of 151bp paired-end reads (SeqCenter) of the resultant strain. Reporter activity was validated using an X-gal (5-bromo-4-chloro-3-indolyl-β-D-galactopyranoside) plate assay by streaking onto a rich defined medium (Chemical Salts Broth, CSB)-agar plates (1x CSB, 100 µg/mL spectinomycin, 1 mM of 2′,4′,6′-trihydroxyacetophenone monohydrate (IPTG), and 80 µg/mL of 2′,4′,6′-

Trihydroxyacetophenone monohydrate (X-gal)).[7]

Reporter activity was quantitatively measured using a modified Miller assay.[8] For each assay, the reporter strain was grown overnight in 1 mL liquid CSB medium supplemented with 100 μg/mL spectinomycin at 37 °C. The resulting cell culture was used to inoculate 1 mL of fresh CSB medium with a 1:50 dilution and supplemented with 1 mM IPTG, 100 μM riboflavin and/or 200 μM **A1** upon initial inoculation. The added ligands were present throughout the outgrowth period (~6 h). Following outgrowth, the cells in 1 mL of each culture were collected by centrifugation at 16,000x*g* for 1 min, washed with liquid CSB medium, then resuspended in 1500 μL of working buffer (60 mM $Na_2HPO_4$, 40 mM $NaH_2PO_4$, 10 mM KCl, 1 mM $MgSO_4$ and 20 mM β-mercaptoethanol). The cells were divided into three separate 500 μL reactions (technical replicates). Using working buffer as a blank, the absorbance at 600 nm for each reaction was measured and subsequently lysed by adding 6.25 μL 15 mg/mL lysozyme and incubating at 37 °C for 20 min. To each reaction, 93.8 μL of 4 mg/mL *ortho*-nitrophenyl-β-D-galactopyranoside (ONPG) was added and allowed to react for 15 min; the reaction was quenched using 250 μL of 1 M $Na_2CO_3$. Absorbances of each solution were measured at 420 nm. For each condition, this process was performed three times, each with a unique isolated colony to yield three biological replicates. Expression was quantified using the equation

$$Miller\ Units = 1000 * \frac{A420}{Reaction\ time\ (minutes) * Reaction\ Volume\ (mL) * A600}$$

where the *A* values represent the absorbance at a specific wavelength.

**General Synthetic Methods**

NMR spectra were collected on a Bruker UltraShield™ NMR spectrometer (600 MHz). Silica gel flash column chromatography was conducted using a Biotage Isolera One purification system. Preparative reversed-phase HPLC was operated on a Waters system (Pump: Waters 1525; Absorbance detector: Waters 2487; Column: Waters Sunfire C18 OBD 5 μm, 19 ×150 mm S-14). The purify was carried out using a linear gradient elution from 0% to 100% methanol (MeOH) in water over 60 min at a flow rate of 5 mL/min. Analytical HPLC was used to assess compound purity on a Waters Symmetry C18 column (5 μm, 4.6 × 150 mm) at a flow rate of 1 mL/min under the same gradient and solvent conditions, with UV absorbance monitored at 254 nm. High-resolution mass spectra (HRMS) were acquired using an Orbitrap Exploris 120 mass spectrometer (Thermo Fisher Scientific) operating in positive electrospray ionization (ESI) mode, coupled to a Vanquish HPLC system (Thermo Fisher Scientific).

**Synthetic route of compound A1**

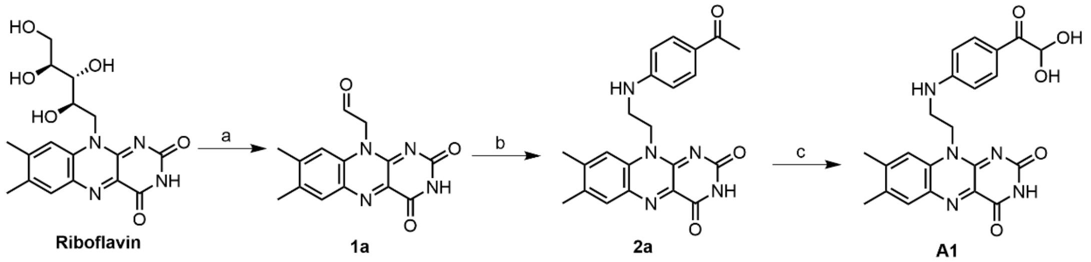


**Reagents and conditions**: **a**. i) $NaIO_4$, $H_2O$, r.t., 12 h; ii) toluene, 110 °C, 5 h; **b**. Sodium cyanoborohydride ($NaCNBH_3$), acetic acid (AcOH), methanol (MeOH), rt., 12 h; **c**. Selenium dioxide ($SeO_2$), 1,4-dioxane/$H_2O$ = 9/1, 100 °C, 24 h.

**2-(7,8-Dimethyl-2,4-dioxo-3,4-dihydrobenzo[*g*]pteridin-10(2*H*)-yl)acetaldehyde** (**1a**)

The preparation of **1a** was performed according to the reported protocol.[9] To a suspension of riboflavin (2.0 g, 5.3 mmol) in water (60 mL), was added sodium periodate (3.4 g, 15.9 mmol). The resulting mixture was stirred at room temperature in the dark for 6 h. The precipitate was filtered off and washed with 50 mL of water for three times, before drying *in vacuo* to yield a yellow solid, which was taken up in toluene (100 mL). The resulting suspension was heated to 110 °C for 5 h, cooled to room temperature, filtered, and dried to obtain compound **1a** as a yellow solid which was used without further purification (1.3 g, 86%). LC−MS: calculated for $C_{14}H_{12}N_4O_3$ $[M + H]^+$ 285.3, found 285.3.

**10-(2-((4-Acetylphenyl)amino)ethyl)-7,8-dimethylbenzo[*g*]pteridine-2,4(3*H*,10*H*)-dione** (**2a**)

To a suspension of **1a** (200 mg, 0.7 mmol) in MeOH (30 mL) was added 1-(4-aminophenyl)ethan-1-one (76 mg, 0.6 mmol) and AcOH (2 mL). The resulting mixture was stirred at room temperature in the dark for 3 h. $NaCNBH_3$ (86.8 mg, 1.4 mmol) was added into the reaction, which was stirred for another 8 h. The reaction solution was filtered, and the resulting yellow solution was concentrated *in vacuo* to yield a yellow residue, which was purified by chromatography on a silica gel column with $CH_2Cl_2$/MeOH to give yellow intermediate **2a** (32 mg, 11%). [1]H NMR (600 MHz, DMSO-$d_6$) δ 11.40 (s, 1H), 7.89 (s, 1H), 7.76 (d, *J* = 8.2 Hz, 2H), 7.52 (s, 1H), 6.70 (d, *J* = 8.1 Hz, 2H), 6.64 (t, *J* = 6.2 Hz, 1H), 4.75 (t, *J* = 6.0 Hz, 2H), 3.64 (dd, *J* = 12.4, 6.1 Hz, 2H), 2.44 (s, 3H), 2.37 (s, 3H), 2.27 (s, 3H). [13]C NMR (150 MHz, DMSO-$d_6$) δ 195.61, 160.47, 156.04, 152.81, 150.76, 146.66, 137.49, 136.16, 134.26, 131.67, 131.38, 131.05, 125.91, 116.74, 111.55, 43.55, 39.16, 26.45, 20.99, 19.21. LC−MS: calculated for $C_{22}H_{21}N_5O_3$ $[M + H]^+$ 404.2, found 404.3.

**10-(2-((4-(2,2-Dihydroxyacetyl)phenyl)amino)ethyl)-7,8-dimethylbenzo[*g*]pteridine-2,4(3*H*,10*H*)-dione** (**A1**)

To a suspension of **2a** (12 mg, 0.03 mmol) in 1,4-dioxane/$H_2O$ = 9/1 (4 mL) was added $SeO_2$ (10 mg, 0.08 mmol). The resulting mixture was heated to 100 °C and stirred for 24h. The reaction mixture was purified by reversed-phase high-performance liquid chromatography (RP-HPLC, MeOH/$H_2O$) to obtain compound **A1** (6 mg, 46%). [1]H NMR (600 MHz, DMSO-$d_6$) δ 11.39 (s, 1H), 7.90 – 7.82 (m, 3H), 7.51 (s, 1H), 6.76 – 6.67 (m, 3H), 6.59 (s, 1H), 5.35 (s, 1H), 4.75 (t, *J* = 6.4 Hz, 2H), 3.68 – 3.61 (m, 2H), 2.37 (s, 3H), 2.26 (s, 3H). [13]C NMR (150 MHz, DMSO-$d_6$) δ 192.12, 160.47, 156.03, 153.25, 150.76, 146.69, 137.48, 136.16, 134.28, 132.30, 131.68, 131.38, 131.34, 122.07, 116.74, 111.49, 96.02, 54.29, 43.54, 39.09, 20.96, 19.19. HRMS: Dicarbonyl form **A1'** was detected, calculated for $C_{22}H_{19}N_5O_4$ $[M + H]^+$ 418.1510, found 418.1510.

Compound **2a**, [1]H NMR, DMSO-$d_6$, 600 MHz.

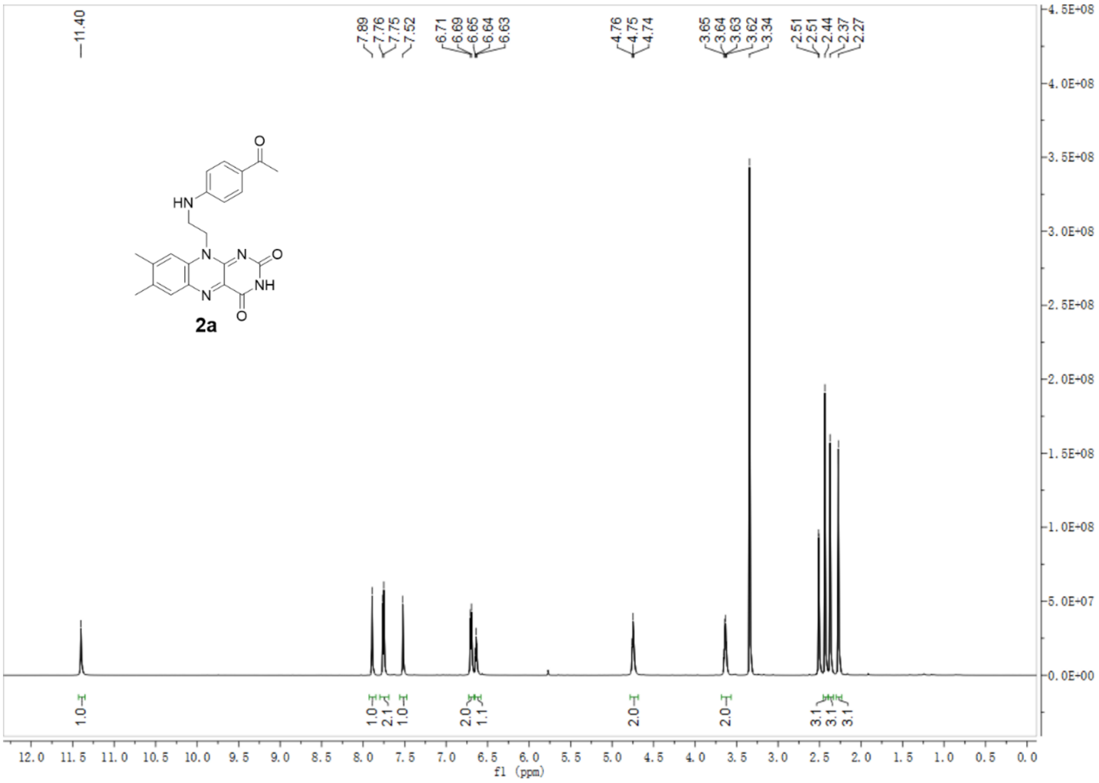


Compound **2a**, [13]C NMR, DMSO-$d_6$, 150 MHz.

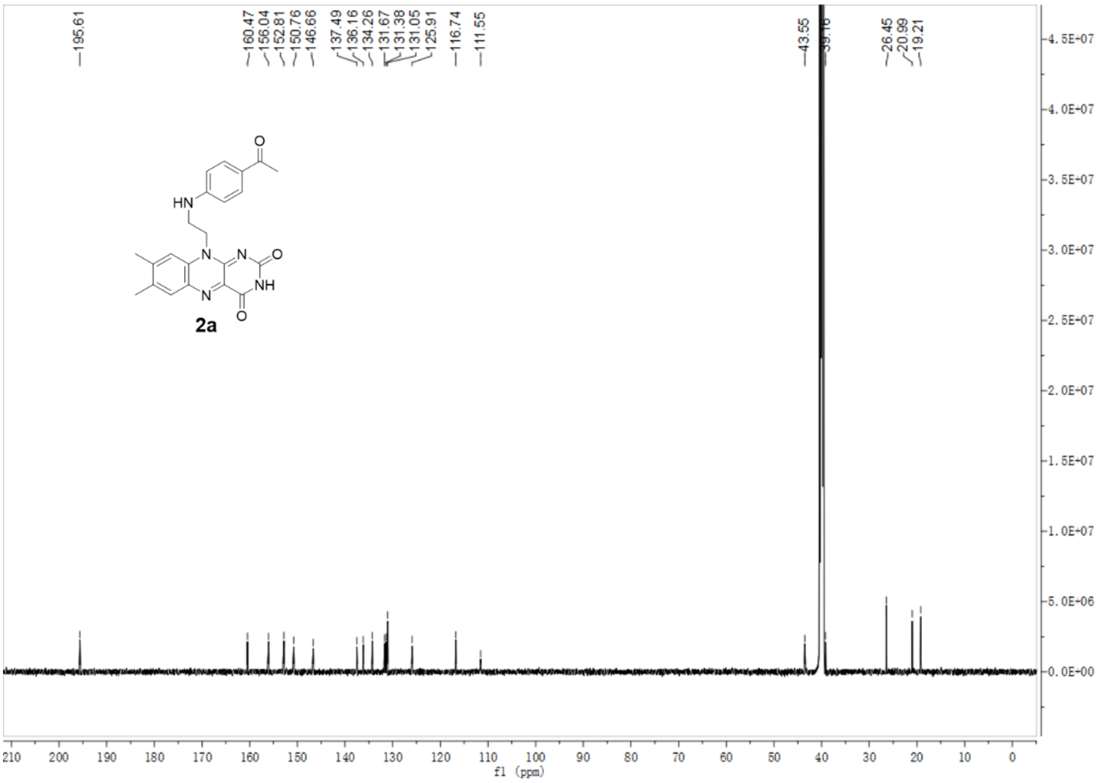


Compound **A1**, $^{1}H$ NMR, DMSO-$d_6$, 600 MHz.

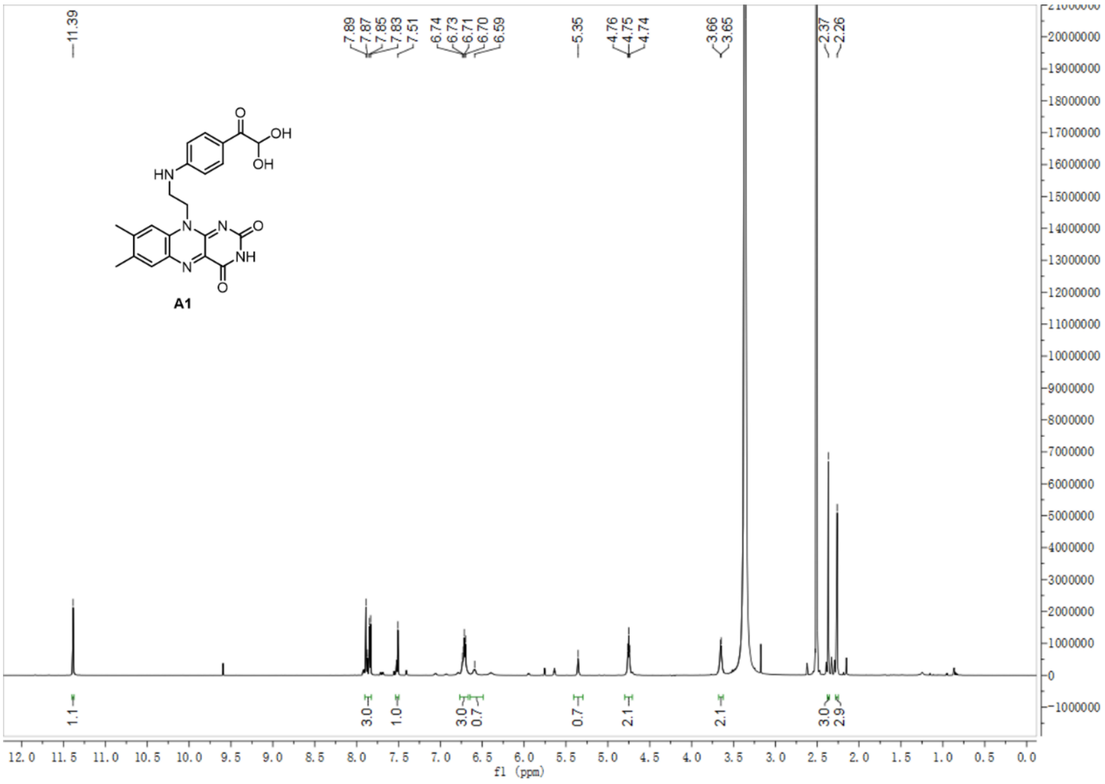


Compound **A1**, $^{13}C$ NMR, DMSO-$d_6$, 150 MHz.

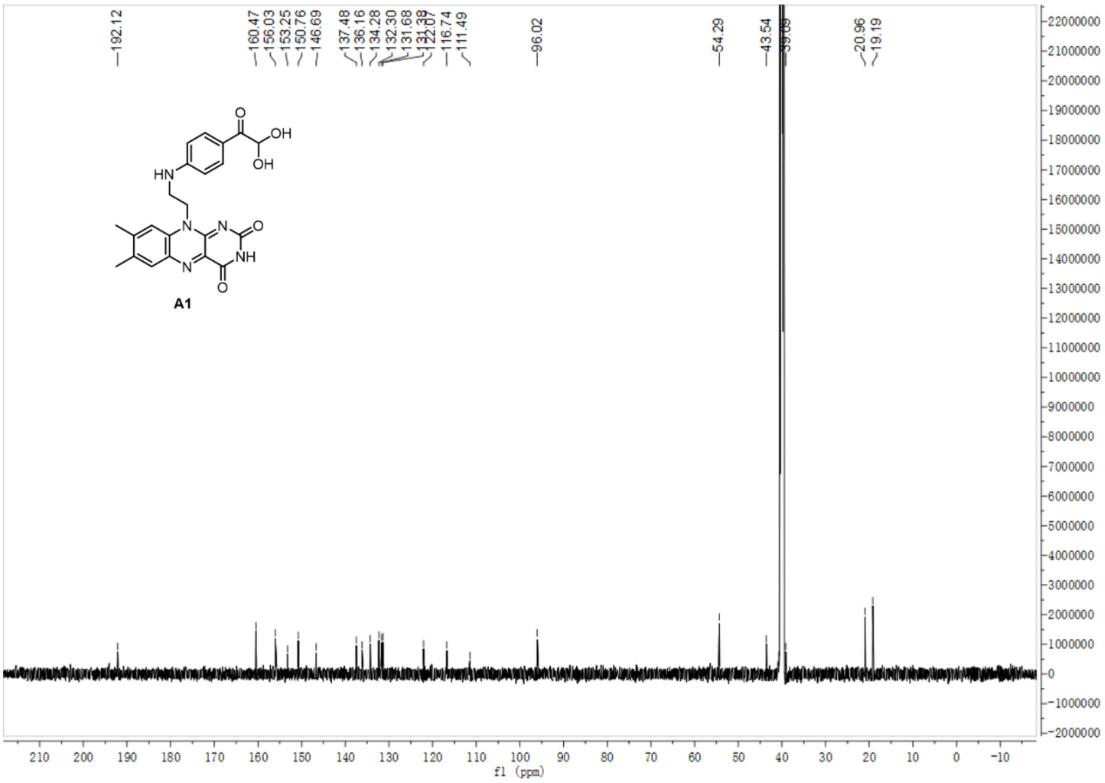


HPLC trace of compound **A1.**

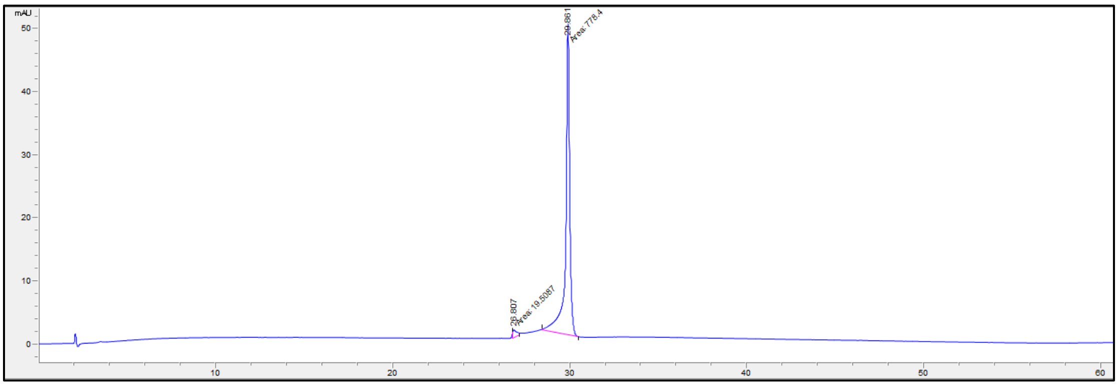


Signal at 255 nm.

| RT (min) | Area | Height | Width | Area % |
|---|---|---|---|---|
| 26.807 | 19.5 | 1.3 | 0.25 | 2.4 |
| 29.861 | 778.4 | 49.3 | 0.26 | 97.6 |

### References

(1) Yang, X.; Wang, J.; Springer, N. A.; Zanon, P. R. A.; Jia, Y.; Su, X.; Disney, M. D. Mapping small molecule-RNA binding sites via Chem-CLIP synergized with capillary electrophoresis and nanopore sequencing. *Nucleic Acids Res* **2025**, *53* (6). DOI: 10.1093/nar/gkaf231.

(2) Springer, N. A.; Zanon, P. R. A.; Taghavi, A.; Sung, K.; Disney, M. D. Discovery of RNA-reactive small molecules guides design of electrophilic modules for RNA-specific covalent binders. *bioRxiv* **2025**. DOI: 10.1101/2025.04.22.649986.

(3) Richard A. Friesner; Jay L. Banks; Robert B. Murphy; Thomas A. Halgren; Jasna J. Klicic; Daniel T. Mainz; Matthew P. Repasky; Eric H. Knoll; Mee Shelley; Jason K. Perry; et al. Glide: A New Approach for Rapid, Accurate Docking and Scoring. 1. Method and Assessment of Docking Accuracy. *J. Med. Chem.* **2004**, *47*, 1739-1749.

(4) Dorado, Oxford Nanopore Technologies. https://github.com/nanoporetech/dorado.

(5) Danecek, P.; Bonfield, J. K.; Liddle, J.; Marshall, J.; Ohan, V.; Pollard, M. O.; Whitwham, A.; Keane, T.; McCarthy, S. A.; Davies, R. M.; et al. Twelve years of SAMtools and BCFtools. *Gigascience* **2021**, *10* (2). DOI: 10.1093/gigascience/giab008.

(6) Wilson, G. A.; Bott, K. F. Nutritional factors influencing the development of competence in the Bacillus subtilis transformation system. *J Bacteriol* **1968**, *95* (4), 1439-1449. DOI: 10.1128/jb.95.4.1439-1449.1968.

(7) Ceres, P.; Garst, A. D.; Marcano-Velázquez, J. G.; Batey, R. T. Modularity of select riboswitch expression platforms enables facile engineering of novel genetic regulatory devices. *ACS Synth Biol* **2013**, *2* (8), 463-472. DOI: 10.1021/sb4000096.

(8) Krute, C. N.; Seawell, N. A.; Bose, J. L. Measuring Staphylococcal Promoter Activities Using a Codon-Optimized β-Galactosidase Reporter. *Methods Mol Biol* **2021**, *2341*, 37-44. DOI: 10.1007/978-1-0716-1550-8_6.

(9) Murahashi, S.; Zhang, D.; Iida, H.; Miyawaki, T.; Uenaka, M.; Murano, K.; Meguro, K. Flavin-catalyzed aerobic oxidation of sulfides and thiols with formic acid/triethylamine. *Chem Commun (Camb)* **2014**, *50* (71), 10295-10298. DOI: 10.1039/c4cc05216a.
